# miRViz: a novel webserver application to visualize and interpret microRNA datasets

**DOI:** 10.1101/813550

**Authors:** Pierre Giroux, Ricky Bhajun, Stéphane Segard, Claire Picquenot, Céline Charavay, Lise Desquilles, Guillaume Pinna, Christophe Ginestier, Josiane Denis, Nadia Cherradi, Laurent Guyon

## Abstract

MicroRNAs (miRNAs) are small non-coding RNAs that are involved in the regulation of major pathways in eukaryotic cells through repression of their target genes at the post-transcriptional level^1^. While high-throughput approaches are broadly used to decipher the biological relevance of miRNAs, extraction of significant information from large miRNA datasets remains challenging. For example, sequencing technologies can quantify the relative expression of up to thousands of mature miRNAs under various experimental conditions. However, in such datasets, small subsets of miRNAs can often show significant differential expression, and deciding which one(s) should be further analyzed can prove difficult. Thus, the current challenge resides in objective analysis, interpretation and visualization of these large datasets, for which specifically suited methods are lacking.

---

Here, we present miRViz (http://mirviz.prabi.fr/), a webserver application designed to visualize and interpret large miRNA datasets, with no need for programming skills. MiRViz has two main goals: (1) to help biologists to raise data-driven hypotheses; and (2) to share miRNA datasets in a straightforward way through publishable quality data representation, with emphasis on relevant groups of miRNAs.

## Results

As summarized in Figure 1, datasets loaded into miRViz can be overlaid onto five proposed miRNA networks (described below), to display graphical representations in which each node corresponds to a given miRNA. MiRViz is designed to work with 11 different species, including human (hsa), mouse (mmu), *C. elegans* (cel) and *Drosophila* (dme). Practical examples, and step-by-step procedures to produce figures are provided as **Supplementary Notes 1-5**.

**Figure 1.**
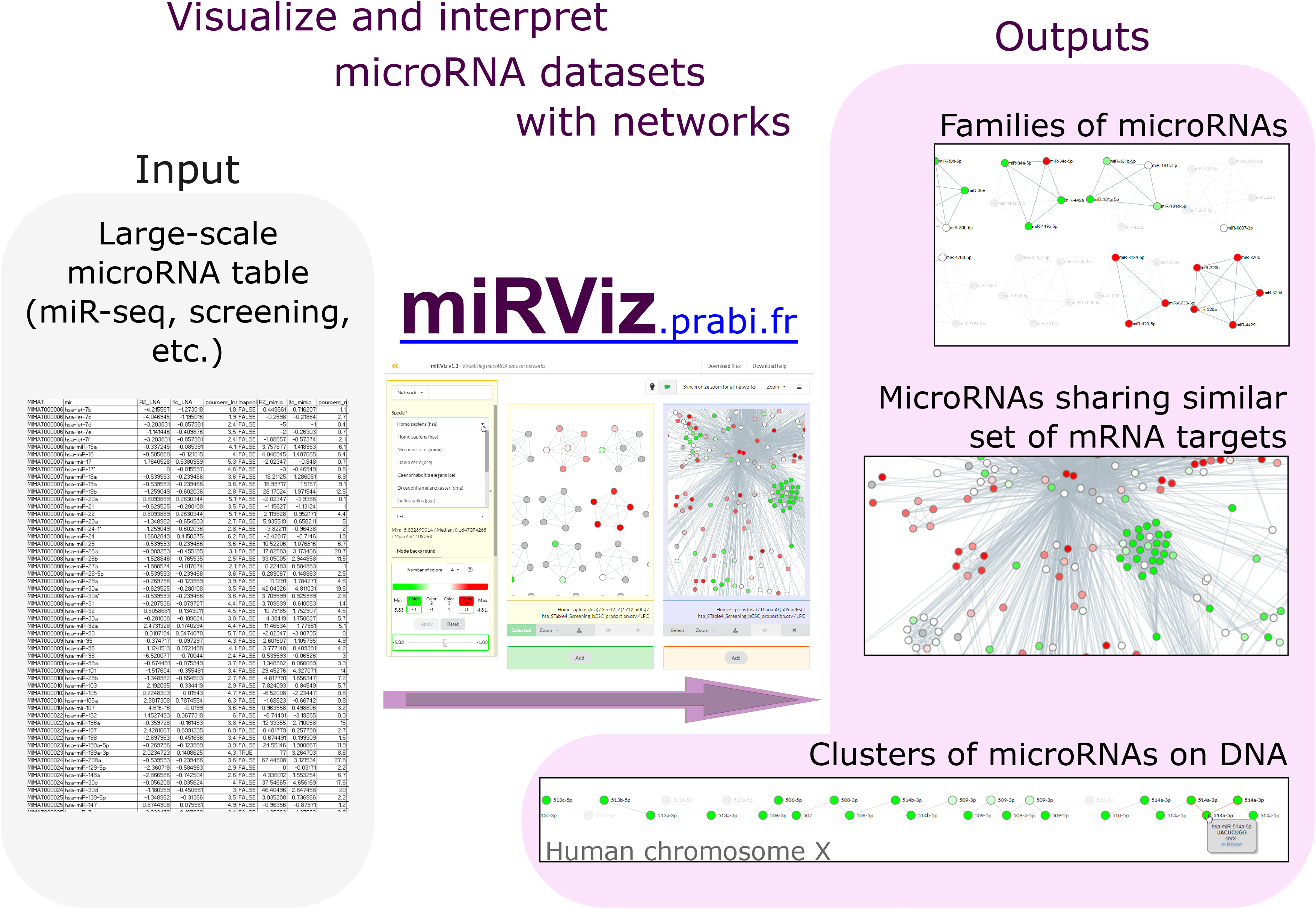
Users load their own datasets in coma-separated values (csv) format through the ‘Load data’ menu, and can choose from among the five different miRNA networks. They can also choose the color code for the nodes, to depict the quantitative values of their data. To compare datasets, up to four windows can be displayed concurrently, and can be manipulated synchronously.

The first network used by miRViz, “Seed2_7”, allows direct visualization of miRNA families by connecting pairs of miRNA nodes that share the same seed sequence. In **Supplementary Figure and Note 1**, we show a practical example of using this network to analyze the cooperative export of the whole hsa-miR-320 family in exosomes^2^.

The two miRViz networks entitled “Genomic_distance” connect neighboring miRNA genes on the genome (**Supplementary Note 2**). **Supplementary Figure 2** shows direct identification and visual representation of the 14q32 cluster of miRNAs with miRViz. This cluster corresponds to loss of heterozygosity of this genomic region for patients with adrenocortical carcinoma with good prognosis^3^. MiRViz enables clear visualization and direct identification of the miRNA cluster.

The last two miRViz networks connect miRNA nodes that share common mRNA targets^4^ (**Supplementary Note 3**). As proof-of-concept, **Supplementary Figure 3** compares two datasets and highlights a cluster of overexpressed miRNAs in pluripotent stem cells^5^, which are active *in vitro* to maintain the equilibrium of breast cancer stem cells^6^.

MiRViz provides various tools for in-depth analysis of miRNA datasets. It might also prove extremely useful in the analysis of data from emerging technologies, like single-cell miRNA-seq experiments, for which proof of concept was recently established^7^, or for profiling rare materials, such as human liquid biopsies.

## Data availability

All of the datasets analyzed during this study are publicly available and are described in detail in **Supplementary Note 4**. In addition, we have described all of the steps to reproduce the Supplementary Figures (**Supplementary Note 5**).

## Code availability

The webserver miRViz, http://mirviz.prabi.fr/, is freely available without login requirements. The source code is available under the Open Source CECILL-B license in the download section.

## Acknowledgements

We thank C. Cochet and JJ. Feige for helpful discussion, and all external testers of miRViz. We thank Christopher Berrie for scientific English editing.

## Author contributions

LG and CC designed the webserver. LG, PG and RB built the miR networks. SS conceived the architecture of the webserver. SS and CP coded the webserver. CC and LG carried out functional tests of the webserver. LG and LD analyzed the miR datasets. GP, CG, JD and NC performed the experiments. LG wrote the manuscript with input and approval from all authors.

