## Supplementary Notes for "miRViz: a novel webserver application to visualize and interpret microRNA datasets"

### INTRODUCTION

MicroRNAs (miRNAs) are non-coding RNAs of around 22 nucleotides that regulate protein-coding gene products at the post-transcriptional level, by directing the RNA-induced silencing complex (RISC) to its mRNA targets. Canonical binding of miRNAs corresponds to almost perfect Watson-Crick pairing of the so-called 'seed' sequence with its mRNA targets, often in the 3' UTR[1]. The seed sequence comprises the six nucleotides at positions 2 to 7 in the 5' region of the mature miRNA. Due to the small number of nucleotides involved in this target recognition, miRNAs lack specificity, and they often have dozens to hundreds of target mRNAs. Groups of miRNAs that share the same seed sequence bind to similar sets of mRNA targets, and are thus classified as miRNA families[1].

The popularization of high-throughput technologies has generated a vast quantity and diversity of large datasets of miRNAs, together with quantification of their various metrics. For example, microarrays and sequencing technologies measure miRNA expression levels under various experimental conditions, to provide data that are often converted into differential expression levels[2–5]. Similarly, phenotypic high-content screening has led to large functional datasets[6]. Users have thus to deal with large tables of quantitative data – usually converted to ‘scores’ – that are arranged as numerical columns. The common questions are then how to interpret these tables, and which miRNAs to select for further evaluation and validation. In the following, we define as ‘hits’ those miRNAs with high scores that indicate their potential interest for biologists. These can be differentially expressed miRNAs, highly expressed miRNAs, or miRNAs with high scores or small p-values.

To help in the analysis of such long lists of miRNAs, we have designed and built a free to use webserver to visualize and interpret miRNA datasets, entitled miRViz (http://mirviz.prabi.fr/). With miRViz, users can visualize their own and/or pre-loaded miRNA datasets on up to five different miRNA networks, with various options to highlight or hide subsets of the data. No operations on the data are proposed through miRViz: such analysis should be conducted before using miRViz. For example, differential expression should be performed with one of the numerous available tools, such as DESeq2[7], before visualization in miRViz. MiRViz is designed to be as intuitive as possible. Additionally, the miRViz help file (‘Download help’, top-right) guides users, click by click.

This Supplementary file is organized as follows. Supplementary Note 1 presents the Seed2_7 network together with a practical example of its use with miRNA-seq expression data. Supplementary Note 2 illustrates the two ‘Genomic_Distance’ networks with miRNA survival data. In Supplementary Note 3, the two networks ‘Diana50’ and ‘TargetScan54’ are presented, together with the comparison of two different datasets using miRViz. The datasets used throughout this study are presented in Supplementary Note 4, and the annotated step-by-step procedures used to produce the Figures shown in this article are described in Supplementary Note 5. Supplementary Tables 1-5 correspond to the Supplementary Figures, and allow their easy reproduction with miRViz.

### NOTE 1. Seed2_7 network

#### a) Description of the Seed2_7 network

For the networks proposed in miRViz, each node is represented by a circle that corresponds to a unique mature miRNA. The seed2_7 networks can be visualized through the miRViz website (http://mirviz.prabi.fr/) for all of the 11 species proposed, with the human version shown in **Supplementary Figure 1A-C**. To build this network, all of the miRNAs were retrieved from the miRBase v22.1 database[8] from a given species, and two nodes were connected if the corresponding miRNAs shared the same seed sequence, as classically defined as nucleotides 2 to 7 from the 5’-end of the mature miRNA. Seed2_7 then displays groups of interconnected nodes, called clusters, which are families of miRNAs that are defined by their common seed sequence. Due to the importance of the seed sequence for mRNA target recognition[1], all of the miRNAs from a given family are believed to share biological functions. Co-expression of miRNAs from the same family thus corresponds to functional redundancy.

To construct the network, the clusters were ranked from largest to smallest, while only considering clusters of two nodes or more. Individual miRNA nodes were removed; *i.e.* the mature miRNAs that do not share their seed sequence with others are removed. In human, the seed2_7 network contains 1,712 miRNAs, and the biggest cluster, which encompasses 27 nodes, corresponds to the hsa-miR-548-5p/559 family with seed sequence AAAGUA. In mouse, the network contains 1133 mature miRNA nodes, and the biggest cluster that contains 14 nodes corresponds to the mmu-let-7-3p/miR-98-3p family of the seed sequence UAUACA. On the website, moving the mouse pointer onto a given node provides information on the corresponding mature miRNA: official name, nucleotide sequence at positions 1-8 from the 5' end (with the seed sequence in bold), chromosome location, and a link to the corresponding miRNA in miRBase[8], http://mirbase.org/.

#### b) MiRNAs of the miR-320 family are selectively exported into colon cancer cell exosomes

As a demonstration of how miRViz can be used to interpret miRNA expression datasets, a few click tutorials are provided in the help file (top-right button on the website), which shows how to visualize tissue-specific miRNAs. Here, we propose a complementary example using the publicly available miR-seq dataset that profiles the LIM1863 colon cancer cell line and three different sorts of extracellular vesicles isolated from the culture supernatant: shed microvesicles, and two types of immunoaffinity-isolated exosomes (i.e., A33, EpCAM)[9]. **Supplementary Figure 1A-C** shows the differentially expressed miRNA families, with the nodes that correspond to the miRNAs with very low expression under all of the conditions set as semi-transparent. MiRNAs with lower and higher expression in vesicles compared to cells are colored in green and red, respectively. Interestingly, miRNAs tend to have similar differential expression in each family, which is highlighted by miRViz. This suggests active export of specific family members through exosomes, as also shown in some experimental cases[10,11]. MiRViz can quickly identify these families, and provide a way to share the result. The hsa-miR-378/422a family is exported specifically in immunoaffinity-isolated A33 exosomes (p-value = 0.008; paired Wilcoxon test; **Supplementary Figure 1A**). The hsa-miR-320 family is significantly exported in both sorts of exosomes (p-value = 0.002; paired Wilcoxon test; **Supplementary Figure 1A, B**). This export is clear only for exosomes (i.e., not for small vesicles), and stronger for the A33 exosomes. We have confirmed these data by independent RT-qPCR measurements (**Supplementary Figure 1D**). Finally, all of the five expressed members of the hsa-let-7-3p/miR-98-3p family are significantly exported in all of the three types of extracellular vesicles (p-value = $6 \times 10^{-5}$; paired Wilcoxon test; **Supplementary Figure 1A-C**).

In some cases, experimental set-ups that measure rare materials impose the possibility of only one experiment per condition with high-throughput techniques, and can validate the results on only a few genes of interest with more sensitive methods. Indeed, Li *et al.* sequenced each condition only once. As a consequence, the classical selection procedure using testing and p-value thresholding is not straightforward. Here, we used miRViz to visually identify differentially expressed families of miRNAs, we performed a test on the whole families of interest, and then we validated this in a low-throughput but sensitive experiment. By extension, we believe that miRViz is particularly interesting for use with rare materials, such as with human or animal samples, or if the signal-to-noise ratio of the data is low, such as in single-cell experiments.

### NOTE 2. Genomic_Distance networks

#### a) Description of the Genomic_Distance networks

Both Genomic_Distance networks can be visualized with the miRViz website (http://mirviz.prabi.fr/) for all of the 11 species, and the human 50k version is shown in

**Supplementary Figure 2B**. To build the networks, we also retrieved all of the miRNAs from the miRBase v22.1 database[8] for a given species, together with their genomic positions. We connected two miRNA nodes to their nearest neighbor if they are closer than a fixed distance apart on the genome. The distance between two connected nodes in pixels is proportional to the logarithm of the chromosomic distance in nucleotides. We gathered 2,883 mature miRNAs for both networks in human. Chromosomes are organized from top to bottom (in human, 1 to 22; X, Y). We fixed the distance threshold at 2 kilobases (kb) for the 'Genomic_Distance_2k' network to visually identify polycistronic miRNA clusters[12]. The 2k human network contains 1,146 links (i.e., pairs of connected nodes). Genomic_Distance networks naturally display groups of connected miRNAs, called clusters, which can correspond to polycistrons. Two miRNA nodes separated by more than 2 kb can be localized in the same cluster step by step. For example, the biggest human cluster contains 26 mature miRNAs, from hsa-miR-376c-5p to miR-655-3p, and it is 10 kb long, although each pair of miRNAs is closer than 2 kb.

The 'Genomic_Distance_50k' is defined with a threshold distance of 50 kb, which leads to larger clusters. It is designed to analyze co-expressed miRNAs following large genomic reorganization events, as encountered in cancers. The largest cluster contains 89 mature miRNAs for the 19q13.4 chromosomic location. It spans 122 kb, from hsa-miR-512-5p to miR-373-5p. This primate-specific cluster is known as C19MC[13].

#### b) High expression of miRNAs of the Xq27.3 cluster is predictive of better prognosis in adrenocortical carcinomas

To demonstrate the interest of the 'Genomic_Distance_50k' network in the context of cancer, we reanalyzed the public data from Assie *et al.*[14], which contains both the miRNome of tumor samples from patients diagnosed with adrenocortical carcinoma and their overall survival (OS) information. For each miRNA, the patients were separated into two groups of equal size, which depended on the miRNA quantification, and a p-value was calculated on the OS after log-rank tests. **Supplementary Figure 2A** shows the Kaplan-Meier curves of two microRNAs of interest, together with the p-value of the log-rank test, and the node colorized according to the p-value. **Supplementary Figure 2B** shows the p-value in a log10 scale overlaid onto Genomic_Distance_50k network, zoomed in on chromosomes 14 to X, with a green gradient for good prognosis miRNAs (miRNAs for which high expression is correlated with good prognosis for the patient), and a red gradient for poor prognosis miRNAs. Two large clusters show up in miRViz:

- cluster 14q32.2 (spanning 197 kb) that is predictive of poor prognosis; i.e., patients who show high expression of the miRNAs of the cluster are associated with shorter OS;
- cluster X27q3 (spanning 95 kb) that is predictive of good prognosis; i.e., patients who show high expression of these miRNAs are associated with longer OS.

While both clusters were described in the original publication[14], miRViz proposes a rapid method to easily identify such clusters and a way to visualize the data. Additionally, in **Supplementary Figure 2C** three clusters are highlighted in blue: hsa-miR-450b-5p/503-5p/424-5p, located in Xq26.3 and associated with adverse prognosis; and two clusters of the hsa-miR-29 family, located in 1q32.2 and 7q32.3, and associated with good prognosis. It is interesting to note that the mature miRNAs from the -3p strand of the miR-29 families that are transcribed from both chromosomes 1 and 7 share the same seed, AGCACC, which suggests redundancy.

MiR-503 has already been described[15], but this is, to the best of our knowledge, the first mention of correlation of the whole cluster of miRNAs. Also, none of the miR-29 members have been described in adrenocortical carcinoma, even if miR-29 members have been shown to be either oncogenic or tumor suppressors in a variety of other cancer types[16]. In adrenocortical carcinoma, high expression of hsa-miR-29b-3p and miR-29a-3p correlate with good prognosis of the patient ($p = 8.2 \times 10^{-5}$ and $5.5 \times 10^{-4}$, respectively, for OS, and $p < 5 \times 10^{-4}$ for disease recurrence). MiR-29 members have more often been described as down-regulated in cancers. Their functional roles as tumor suppressors have been associated with cancer progression restriction by promotion of tumor-cell apoptosis, with suppression of DNA methylation of tumor-suppressor genes, and inhibition of tumor-cell proliferation, as well as increased chemosensitivity[16].

### NOTE 3. Co-regulation networks (Diana50 and TargetScan54)

#### a) Description of Diana50 and TargetScan54 networks

Both ‘Diana50’, and ‘TargetScan54’, which we call co-regulation networks, can be visualized with miRViz, as in **Supplementary Figure 3**. For these networks, we connected the miRNA nodes if they shared >50% or >54%, respectively, common mRNA targets, as predicted by Diana MicroT v3[17] or Target-Scan v6.2[18]. ‘TargetScan54’ contains 1,530 nodes, each of which corresponds to a mature miRNA. ‘Diana50’ comprises 539 nodes, and gathers fewer but more commonly found miRNAs, and is more easily viewable. We have previously described both networks[19], and in particular, the choice of the thresholds. Briefly, these thresholds of 50% and 54% maximized the ‘betweenness centrality’ measurements of each network, and so reveal miRNA communities. The networks layout was performed using Cytoscape with the unweighted spring embedded layout[19,20]. Nodes are then closer when they form dense clusters of interconnected nodes. However, these clusters were too dense, with overlapping nodes. To facilitate visualization of these clusters, and to better highlight them, we applied the layout again on these clusters separately, and pulled them to the side of the networks. Dense clusters correspond to large groups of miRNAs that share many predicted mRNA targets. As in the example below, co-expression of the miRNAs of a given cluster suggests redundancy in mRNA repression. Thus, functional validation should be performed by simultaneously knocking-down co-expressed miRNAs of a cluster, as individual miRNA knock-down might lead to only subtle phenotypic effects, if any.

#### b) MiRViz visually identifies the miR-302/519 stem-cell family in the regulation of breast cancer stem cell equilibrium

As proof of purpose, **Supplementary Figure 3A** shows the differential expression of miRNAs in stem cells cultured under two different conditions that favor either pluripotency or differentiation. Here, a ‘stem cell’ miRNA cluster clearly shows up in red, which highlights the overexpressed miRNAs in pluripotent stem cells. Most of these miRNAs have already been hypothesized to cooperatively regulate pluripotency[21]. The group comprises miR-17/20/93/106/302/../519/520 with shifted seed sequences (AAAGUG, AAGUGC, AGUGCU), and miR-411 with seed sequence AGUAGA. To determine the efficiency of miRViz to compare different datasets and the possibility that it can raise biological questions of interest, we can compare **Supplementary Figure 3A** and **3B. Supplementary Figure 3B** represents a functional screening dataset where we measured the relative levels of breast cancer stem cells (bCSC) in a human breast adenocarcinoma cell line (SUM159 cells) upon miRNA systematic and individual overexpression[22]. Green (resp. red) nodes represent miRNAs that upon overexpression lead to smaller (resp. higher) bCSC proportions. In our previous study, we focused on the modes of action of miR-600. Here, the miRViz representation highlights the redundant action of the 'miRNA stem cell' cluster on the balance of the bCSC phenotype. It is, however, surprising that miRNAs for which expression was correlated with pluripotency in normal cells indeed lead to decreased proportions of bCSCs when overexpressed (**Supplementary Figure 3B**, highlighted green cluster). This suggests that the fine-tuning of this specific group of miRNAs might have an important and yet unknown role in the maintenance of the 'stem' state of normal and cancer cells. Interestingly, miRViz identifies this group of miRNAs, and suggests target gene redundancy, which might explain why their individual knock-downs in separate experiments (data not shown) had little or no effects on the bCSC equilibrium. It also emphasizes the need to knock-down these miRNAs collectively to restore the expression of the target genes that are responsible for the stem-cell features.

#### c) Diana50 and TargetScan54 structures are correlated with biological functions

We performed gene ontology enrichment on the predicted mRNA targets for each individual miRNA. For a given ontology and miRNA, a small p-value (typically $<10^{-5}$) suggests that the miRNA regulates the corresponding function under certain cellular conditions. **Supplementary Figures 4** and **5** show that miRNAs that are assumed to regulate a given ontology (i.e., pathway or function) are not randomly spread out in the networks. Supporting our previous study[19], the Diana50 and TargetScan54 networks are structured in two parts. The upper part of both networks contains miRNAs that are almost all predicted to regulate gene expression, together with the two more central subnetworks of let-7 and miR-17/93 (**Supplementary Figures 4A, 5A**). The lower parts of both of these networks contain many miRNAs that are predicted to regulate signal transduction through small GTPases (**Supplementary Figures 4B, 5B**). Altogether, these observations show that the Diana50 and TargetScan54 structures correlate with biological predictions, and that the positions of hits in the networks inform the users of the regulated pathways; e.g., miRNA hits in the upper part might be important regulators of gene expression.

### NOTE 4. Datasets. Material and Methods

#### a) Pre-loaded datasets

Three miRNA tables are pre-loaded to ease comparisons with user datasets: 'hsa_miRmine_cells', 'hsa_miRmine_tissues', and 'hsa_TissueAtlas'. For the first two datasets, the data were gathered from miRmine[5]. For the third dataset, the data were gathered from TissueAtlas[4]. MiRNA expression was transformed into log2 scales, and then averaged across all of the experiments performed under the same conditions (the number of different experiments used to average is indicated in brackets). Expression on a log2 scale spans from 0 to 20.

#### b) Large-scale datasets used in the Supplementary Figures

All of the data described below were analyzed with R version 3.4[23]. They are provided in csv format directly reusable in miRViz to reproduce the figures provided here.

#### *MiRNA sequencing of small vesicles and colon cancer cells*

We used colon cancer cells and the exosome miRNA sequencing public dataset SRA106214 produced by Ji and coworkers[9]. The raw data were filtered using FastQC v0.72 (http://www.bioinformatics.babraham.ac.uk/projects/fastqc/) and Trim Galore! v0.4.3.1 (https://www.bioinformatics.babraham.ac.uk/projects/trim_galore/), and aligned against the miRBase v22 sequence list[8] using Bowtie1 v1.2.0[24], with the read counts normalized using DESeq2 v2.11.40.3 [25]. Differential expression is expressed in the log2 scale (log2[miR_count(Exo)/miR_count(Cell)]) in **Supplementary Table 1**.

#### *Adrenocortical cancer survival data*

MiRNA sequencing data from individual patient tumors diagnosed as adrenocortical carcinoma were retrieved from the GSE49279 public dataset. Clinical information, including follow-up and overall survival of the same 45 patients, were retrieved from the Supplementary Materials of Assie *et al.*[14]. For each miRNA, the population of patients was separated into two equal groups of 22 patients by the median expression of the miRNA. P-values were calculated with log rank tests for OS with the R package 'survival' version 2.44. A score was defined by log10 transformation of the p-value, and a minus sign was added depending on the prognosis for patients with high expression of the miRNA (**Supplementary Table 2**). For example, miR-29a-3p has a score of -3.3, which corresponded to a p-value of $5.5 \times 10^{-4}$, with its high expression correlated with good patient prognosis. The node corresponding to this miRNA is pure green in **Supplementary Figure 2**.

#### *MiRNA expression with microarrays in human embryonic stem cells*

The miRNA expression in human embryonic stem cells originates from the public GSE14473 dataset[26]. In this study, 10 different cell lines were separately cultured under conditions to either maintain an undifferentiated state or to promote undirected differentiation. For each miRNA, we took the median among all of the cell lines of the log2 fold-change (LFC) between the undifferentiated and differentiated states (**Supplementary Table 3**).

#### *High-content screening of breast cancer stem cell equilibrium upon miRNA overexpression*

To demonstrate the miRViz use for high-content screening datasets, we used an in-house dataset detailed earlier[22]. Briefly, after the individual transfection of miRNA mimics from a human miRNome-wide library, high-content screening was used to measure the proportion of breast cancer stem cells (ALDEFLUOR-positive cells) in LFC compared to the control (**Supplementary Table 4**). For example, hsa-miR-512-3p has a LFC of -1.1, which means that there were almost half the bCSCs after miR-512-3p transfection compared to control ($2^{-1.1} = 0.46$). This miRNA is represented in light green in **Supplementary Figure 3B**.

#### *Gene ontology enrichment of mRNA targets of individual miRNAs*

For each mature miRNA, the pool of predicted mRNA targets by TargetScan v6.2[27] was retrieved. Gene ontology enrichment was performed on each pool, which provided a p-value for each miRNA and ontology. The p-values were expressed in –log10 scale: a value of 12 (shown in red in **Supplementary Figures 4** and **5**) corresponds to $p = 10^{-12}$ (**Supplementary Table 5**). Ontologies were retrieved using the R package GO.db version 3.1[28], and the correspondence between gene ID databases was performed with biomaRt R package version 2.26[29]. Fisher exact tests were performed to determine the p-values.

#### c) RT-qPCR validation experiments

The colorectal-cancer-derived LIM1863 cell line was purchased from Public Health England (Salisbury, UK). The cells were cultured in RPMI 1640 medium containing 2 mM L-glutamine, 25 mM HEPES, 10% fetal calf serum, 0.6 µg/mL insulin, 1 µg/mL hydrocortisone and 10 µM 1-thioglycerol. The cells were grown as floating organoids in a humidified incubator at 37 °C in a 5% $CO_2$–95% air atmosphere.

Exosomes were prepared using differential centrifugation of conditioned medium collected from LIM1863 cells. Successive centrifugations at increasing speeds were performed to eliminate dead cells and large cell debris (10 min at 300× *g*; 10 min at 2,000× *g*). The supernatant was centrifuged for 30 min at 10,000x *g* at 4 °C, to pellet the microvesicles. The final supernatant was filtered through a Millipore Stericup filtration system (pore size, 0.22 µm), and then ultracentrifuged at 100,000× *g* for 180 min to pellet the exosomes. The pellet was washed with 8 mL phosphate-buffered saline (Gibco, Life Technologies) and ultracentrifuged again at 100,000× *g* for 60 min to eliminate any contaminating proteins. The microvesicles and exosomes were resuspended in 200 µL phosphate-buffered saline and stored at -80 °C.

Total RNA from LIM cells, cell-derived microvesicles, and cell-derived exosomes was isolated using mirVANA PARIS kits (Applied Biosystems, Thermo Fisher Scientific), according to the manufacturer instructions. MiRNA levels were measured using RT-qPCR with TaqMan miRNA assays (Applied Biosystems, Thermo Fisher Scientific). Ten nanograms of total RNA were reverse transcribed using TaqMan miRNA Reverse Transcription kits and miRNA-specific stem-loop primers (Applied Biosystems, Thermo Fisher Scientific) in a 15 µL reverse transcription reaction (composed of 0.15 µL 100 mM dNTPs mix, 1.5 µL 10× reverse transcription buffer, 0.19 µL RNase inhibitor [20 U/µL], 4.16 µL $H_2O$, 1 µL multiscribe reverse transcriptase, and 5 µL input RNA), using a TGradient thermal cycler (Biometra, Goettingen, Germany) at 16 °C for 30 min, 42 °C for 30 min, and 85 °C for 5 min. Real-time PCR was performed on the 5'-extended cDNA with TaqMan 2× Universal PCR Master Mix and the appropriate TaqMan MicroRNA Assay Mix for each miRNA of interest. Briefly, 4.5 µL 2.5-fold diluted reverse-transcribed product was combined with 5.5 µL PCR assay reagents to generate a PCR volume of 10 µL. Real-time PCR was carried out on a C1000 thermal cycler (CFX96 Real-Time system; Bio-Rad) at 95 °C for 10 min, followed by 40 cycles at 95 °C for 15 s and 60 °C for 1 min. The data were analyzed with CFX Manager software version V1.5.534.0511 (Bio-Rad). MiR-16 was used as an endogenous control for normalization. Normalized expression was calculated using the comparative Ct method, and fold-changes were derived from the $2^{-\Delta\Delta Ct}$ values for each miRNA.

### NOTE 5. Step-by-step procedure to reproduce the Figures of this paper

Each Supplementary Figure of the present study can be reproduced easily. First, the corresponding dataset (**Supplementary Tables 1-5**) has to be loaded into the webserver (main menu top left, click on 'Load data'). The downloadable file called 'mirviz-help.pdf' (top right) describes in detail the procedure in a dedicated section.

#### a) Supplementary Figure 1: Differential expression and miRNA families

1. **Load** the 'STable1_Sequencing_Exosome_SRA106214.csv' file into miRViz.

a. MiRNA column should be ‘MIMAT’.
b. Data column should be the last four (‘LFC…’ and ‘Max_Expr’).

2. Back in the ‘Network’ menu, **select the STable1** and LFC_C_vs_A33 in the ‘Colors’ section.
   a. Select the four-color scale to color the nodes. For differential expression, we often advise to set-up for each of the four colors: -2 / -0.5 / 0.5 / 2, and to keep the default colors. If so, strongly repressed miRNAs (LFC <-2) will be associated with a pure green color, moderately repressed miRNAs with a gradient of green (LFC -2 to -0.5), little to no differential expression will be white (LFC -0.5 to 0.5), moderately overexpressed miRNAs with a red gradient (LFC 0.5 to 2), and strongly overexpressed miRNAs pure red (LFC >2).
   b. Grey nodes correspond to unavailable data (miRNAs with no measurement).
3. **Add a new graph** in a second window (the color of the left band will change according to the window selected).
   a. Repeat the same procedure as in section 2 above for the LFC_C_vs_EpCam dataset.
   b. In a third window, repeat the procedure for the LFC_C_vs_sMV dataset.
   c. Three windows should be obtained, with three different datasets to be compared.
   d. Each window can be selected by the ‘select’ button below, or by zooming or moving the network inside a given window.
4. **Hide** miRNAs with very low expression.
   a. Poorly expressed miRNAs with differential expression are often affected by experimental noise. We suggest to hide them to focus on more interesting miRNAs.
   b. Click ‘Display/Hide’ in the top left menu. Select ‘Max_Expr’ from the data column. Select 5 and 21.24 (button pushed to the right), so that only miRNAs with a log2 expression of ≥5 under at least one of the conditions will be displayed.
   c. In Supplementary Figure 1A-C, we chose the semi-transparent option to show the underlying network. When looking for interesting areas, we suggest the ‘transparent’ option.
   d. Alternatively, for differential expression, transparency can be set-up with a p-value (only miRNAs associated with a low enough p-value will appear).
5. **Navigate** the networks to identify groups of interesting miRNAs and compare the datasets.
   a. The top right button ‘Synchronize zoom for all networks allows navigation in the three windows in parallel. Click and move for displacement, and roll the mouse to zoom in or out. Pre-defined zoom levels are also proposed.
   b. Zooming in lets the miRNA name appear near the nodes. Alternatively, the ‘Zoom dependent’ button in the left band can be unclicked, and press ‘s’ to show or remove the miRNA legend.
   c. A high resolution screen is highly recommended to benefit from miRViz. To further increase the size, we recommend full screen size of the browser (F11 or Control+Cmd+F, in most browsers), and to hide the left band of miRViz (double yellow arrows, top left).
   d. To scroll down the whole website, the mouse pointer should be located between two windows (to avoid zooming inside a window).

e. The web browser can be zoomed out to see all three windows at once (often ctrl + roll, with the mouse pointer between two windows).

#### b) Supplementary Figure 2: Survival data and genomic positions

The browser can be refreshed to start a completely new analysis. Alternatively, new data can be added to the previous analysis, or a new browser tab or window can be launched. A lot of loaded data slows down miRViz, as also for many tabs in the browser.

1. **Load** the 'STable2_Survival_AdrenocorticalCarcinoma_GSE49279.csv' file into miRViz.
   a. MiRNA column should be 'miR'.
   b. Data column should be the last two ('Median_expression' and 'log10_OS_sign').
   c. A few rows are ignored as they were not identified with miRViz nodes. Conversion to MIMAT identifiers would minimize the issue. MiRNAs removed from miRBase version 22 or earlier will also not be identified.
2. Change the **network.**
   a. Select the 'Genomic_Distance_50k' network.
   b. Click the 'One column' button, top right (near the 'Zoom' and 'Tool tip' buttons), to enlarge the window of interest.
3. Back in the 'Network' menu, **select the STable2** and log10_OS_sign in the section 'Colors'.
   a. To display p-values, we recommend the log scale. Here, a sign is added in the dataset to separate bad *versus* good prognosis miRNAs. MiRNAs with good prognosis have a negative sign.
   b. Select the four-color scale to color the nodes. Set up: -3 / -1 / 1 / 3, and keep the default colors. MiRNAs associated with OS with a p-value <$10^{-3}$ will be in pure color, as green for good prognosis, and red for bad prognosis. MiRNAs with a p-value of ≥0.1 will be pure white. MiRNAs with intermediate p-value will have a shaded color, as green or red, depending on the prognosis.
   c. Grey nodes correspond to unavailable data (miRNAs with no measurement).
4. **Hide** miRNAs with very low expression.
   a. Poorly expressed miRNAs in most of the patients are often affected by experimental noise. We suggest to hide these to focus on more interesting miRNAs.
   b. Click 'Display/Hide' in the top left menu. Select 'Median_expression' from the data column. Select 10 and 1609091 (button pushed to the right), so that only miRNAs with sufficient expression in at least half of the patients will be displayed.
   c. In Supplementary Figure 2, we chose the semi-transparent option to show the underlying network. When looking for interesting areas, we suggest the 'transparent' option.
5. **Navigation** advice is the same as the previous case with differential expression.

#### c) Supplementary Figure 3A: Differential expression and 'Diana50'

1. **Load** the 'STable3_MicroArray_DE_totipotent_GSE14473.csv' file in miRViz.
   a. MiRNA column should be 'miR'.
   b. Data column should be the last one ('Median_LFC').

c. A few rows are ignored as they were not identified with miRViz nodes. Conversion to MIMAT identifiers would minimize the issue. MiRNAs removed from miRBase version 22 or earlier will also not be identified.

2. Select the **'Diana50' network**.
3. Back in the 'Network' menu, **select the STable3** and Median_LFC in the 'Colors' section.
   a. Select the four-color scale to color the nodes. Set up: -2 / -0.5 / 0.5 / 2 (same as for case 1).
   b. Grey nodes correspond to unavailable data (miRNAs with no measurement).
4. **Hide** miRNAs not measured.
   a. Click 'Display/Hide' in the top left menu. Select 'Median_LFC' from the data column. Push both buttons respectively to the left and right.
   b. In Supplementary Figure 3, we chose the semi-transparent option to show the underlying network. When looking for interesting areas, we suggest the 'transparent' option.
5. **Navigation** advice is the same as in case 1.

#### d) Supplementary Figure 3B: Functional score and 'Diana50'

1. **Load** the 'STable4_Screening_bCSC_proportion.csv' file into miRViz.
   a. MiRNA column should be 'MIMAT'.
   b. Data column should be the last one ('LFC').
   c. A few rows are ignored as they were not identified with miRViz nodes. This is due to the removal of these identifiers in miRBase v22 or earlier.
2. Select the **'Diana50' network**.
3. Back in the 'Network' menu, **select the STable4** and LFC in the 'Colors' section.
   a. Select the four-color scale to color the nodes. Set up: -3 / -1 / 1 / 3 (scale to highlight stronger effects as compared to the scale chosen in cases 1 and 3).
   b. Grey nodes correspond to unavailable data (miRNAs with no measurement).
4. **Hide** miRNAs not measured
   a. Click 'Display/Hide' in the top left menu. Select 'LFC' from the data column. Push both buttons respectively to the left and right.
   b. In Supplementary Figure 3, we chose the semi-transparent option to show the underlying network. When looking for interesting areas, we suggest the 'transparent' option.
5. **Navigation** advice is the same as in case 1.

#### e) Supplementary Figures 4 and 5: Ontology enrichment and co-regulation networks

1. **Load** the 'STable5_GOenrichment_mRNA_targets.csv' file into miRViz.
   a. MiRNA column should be 'MIMAT'.
   b. Data column should be the last two ('Reg. gene expr.' and 'small GTPase mediated sign. transd.').
2. Select the **'Diana50' (Supplementary Figure 4)** or the **'TargetScan54' (Supplementary Figure 5) network**.
3. Back in the 'Network' menu, **select the STable5** and 'Reg. gene expr.' in the section 'Colors'.
   a. Select the **three-color scale** to color the nodes. Set up: Unchanged / 5 / 10. The first color should be turned to white. Thus, the miRNA nodes with p-

values >$10^{-5}$ are in white, those between $10^{-5}$ and $10^{-10}$ are red shaded, and those <$10^{-10}$ are red.

4. **Add a new graph** in a second window (the color of the left band will change according to the selected window).
    a. Repeat the same procedure as section 3 above for the 'small GTPase mediated sign. transd.' Dataset.
5. **Navigation** advice is the same as in case 1.

#### f) Generating high-quality images

Typically, three possibilities are proposed to generate high-quality images of the colored miRNA networks:

1. Intermediate quality, but easy option.
    a. Export as png (export button below each window), and assemble the figure.
    b. Higher resolution images are obtained from higher resolution screens.
2. High quality images option.
    a. Export as pdf – the image is a scalable vector graphic, so that it can be zoomed in without quality loss.
    b. This solution requires high RAM memory, especially with the Genomic_Distance and TargetScan54 networks.
    c. Assemble the figures with vector drawing software (such as Inkscape or CorelDraw).
3. High quality and more flexible option.
    a. Download the cytoscape[20] file through the 'Download file' icon, top right.
    b. Benefit from the flexibility of Cytoscape to adapt your drawings. For example, multiple values can be displayed on each node using the enhancedGraphics plugin[30].

### LEGENDS FOR SUPPLEMENTARY FIGURES

**Supplementary Figure 1. A-C.** Heat map of the differentially expressed miRNAs between the exosomes or microvesicles and the parental LIM1863 cells overlaid on the "Seed2_7" network of miRViz. MiRNA families naturally appear from the largest to the smallest. Red and green nodes correspond to miRNAs overexpressed and repressed in vesicles, respectively. Three interesting clusters are zoomed in on at the bottom right: The miRNA cluster in the red square corresponds to the miR-320 family, the purple hexagon corresponds to the miR-378/422a family, and the blue circle to the let-7-3p/miR-98-3p family. Nodes corresponding to miRNAs not expressed in this cell type were set to semi-transparent. **A.** MicroRNAs in A33 exosomes derived from colon cancer cells *versus* parental cells. **B.** MicroRNAs in EpCAM exosomes derived from colon cancer cells *versus* parental cells. **C.** MicroRNAs in shed microvesicles *versus* parental cells. **D.** Experimental validation using RT-qPCR, showing selective enrichment of the miR-320 family members in the exosomes compared to the parental cells and microvesicles.

**Supplementary Figure 2.** Prognostic potential of miRNAs for overall survival of patients with adrenocortical carcinoma. **A.** Kaplan-Meier curves for miR-514a-5p (top) and miR-411-5p (bottom). On the right are the corresponding nodes colorized using the p-value of the log-rank test and the color scale chosen in **B-C**. **B.** Prognostic value of individual miRNAs overlaid on the 'Genomic Distance 50k' network. Bottom: View of the whole Genomic_Distance_50k network. The square correspond to the zoomed in area displayed above. Chromosomes are organized from top to bottom (1 to 22, X, Y). MiRNAs for which high expression correlates with poor prognosis are highlighted in red. Good prognosis miRNAs are represented in green. MiRNAs with low expression are set as transparent. **C.** MiRViz screen shots of interesting areas that show miRNA names and the action of the mouse pointer on a given node. The squares on the full network below correspond to the interesting areas. MiRNAs with low expression are set as semi-transparent. A few small clusters of miRNAs with high differential expression are highlighted (blue squares): Clusters 1 and 2 correspond to miR-29 family located on chromosomes 1 and 7, and cluster 3 correspond to miR-503-5p/424-5p located on chromosome X. The two major clusters in green and red squares (i.e., Xq27, 14q32) of 95 and 197 kilobases, respectively, show groups of miRNAs associated with good and poor prognosis, respectively.

**Supplementary Figure 3. A.** Differential expression of miRNAs from cells grown in totipotent medium *versus* differentiation medium, as obtained from the GSE14473 public dataset[26], overlaid on the 'Diana50' network. MiRNA nodes in red correspond to miRNAs overexpressed in totipotent cells. **B.** Changes in the bCSC relative proportions after miRNA overexpression. MiRNA nodes in green correspond to miRNAs for which overexpression leads to decreased proportions of bCSCs. **A, B.** Blue squares show the clusters described in the main text, which are zoomed in on at the side of the whole network.

**Supplementary Figure 4.** Gene ontology enrichment for predicted targets of individual miRNAs overlaid on top of the Diana50 network. Red nodes correspond to miRNAs predicted to regulate many protein coding genes known to participate in the following ontologies: (**A**) GO:0010468 (regulation of gene expression); (**B**) GO:0007264 (small-GTPase-mediated signal transduction).

**Supplementary Figure 5:** Same as Supplementary Figure 4, but using the 'TargetScan54' network.
