## Supplementary Figures for "miRViz: a novel webserver application to visualize and interpret microRNA datasets"

### Supplementary Figure 1A

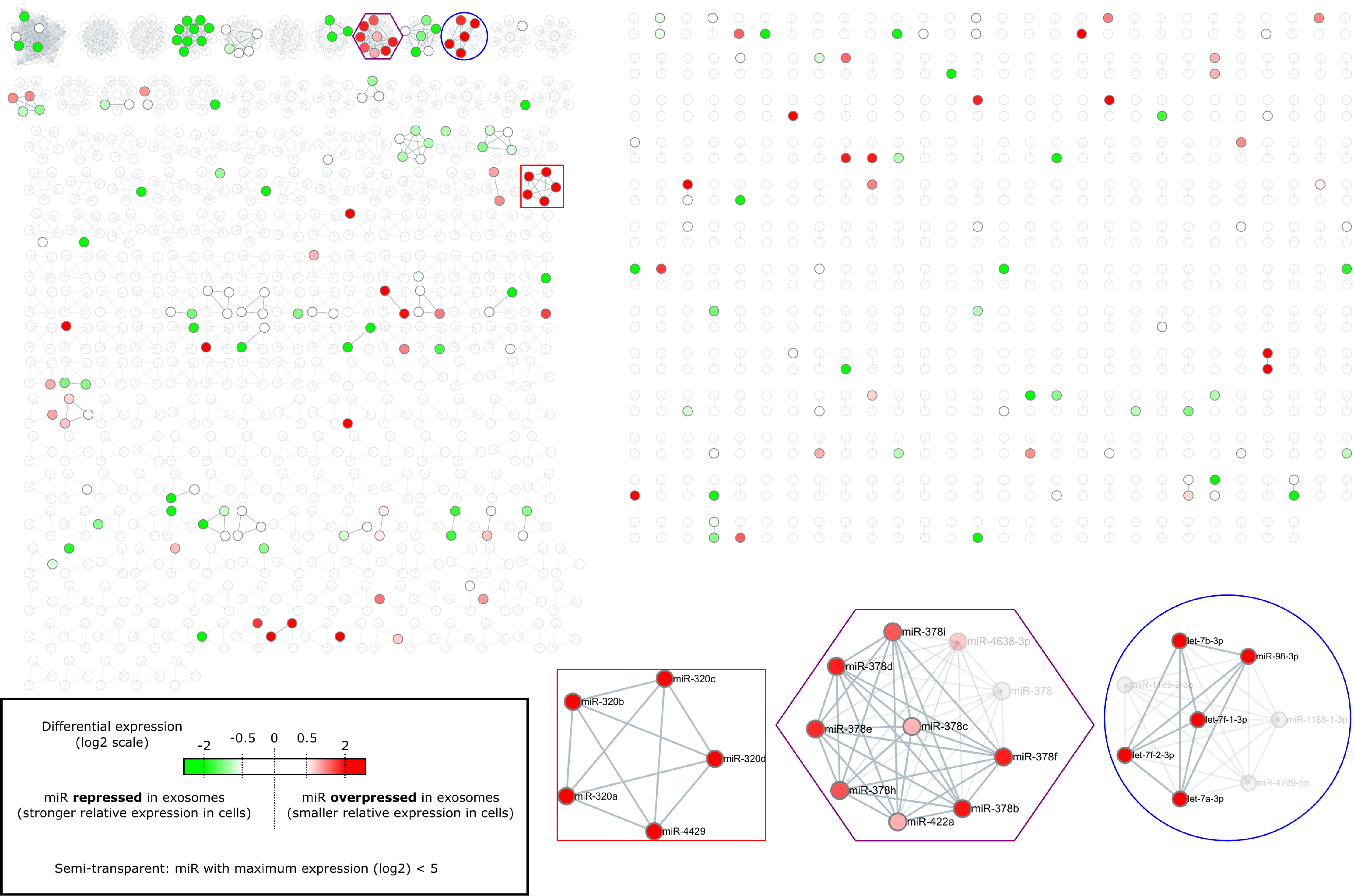

### Supplementary Figure 1B

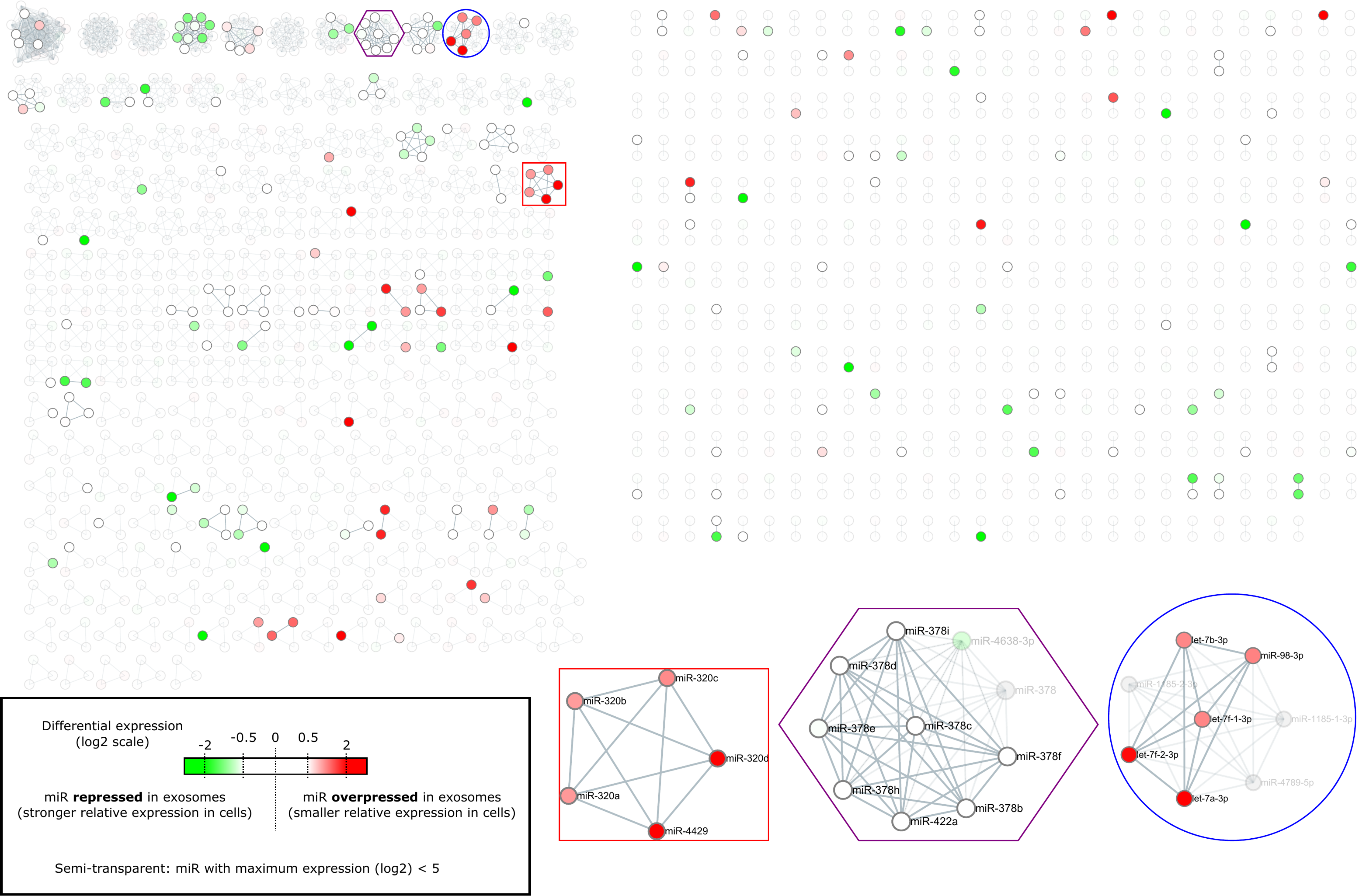

### Supplementary Figure 1C

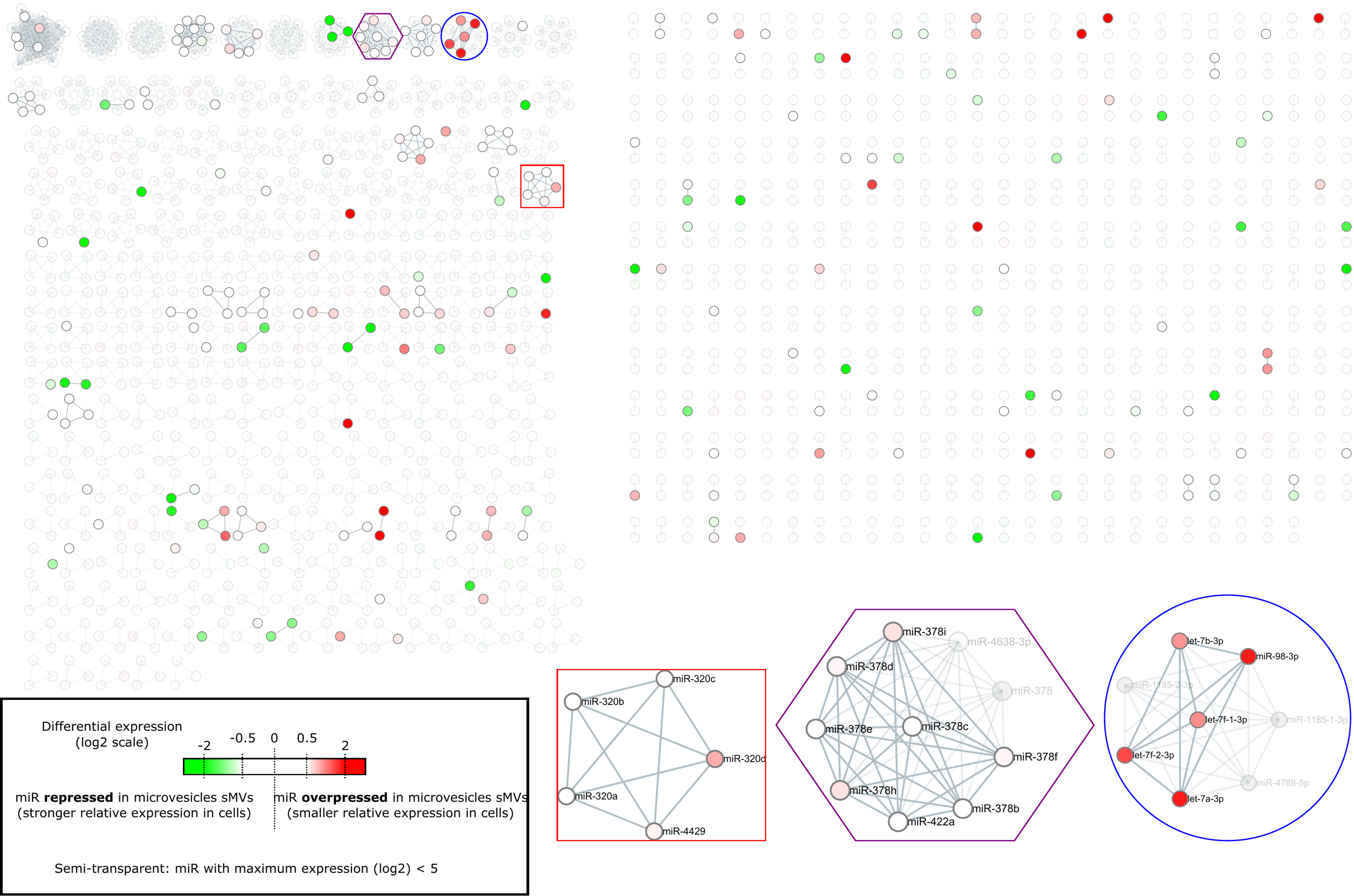

Supplementary Figure 1D

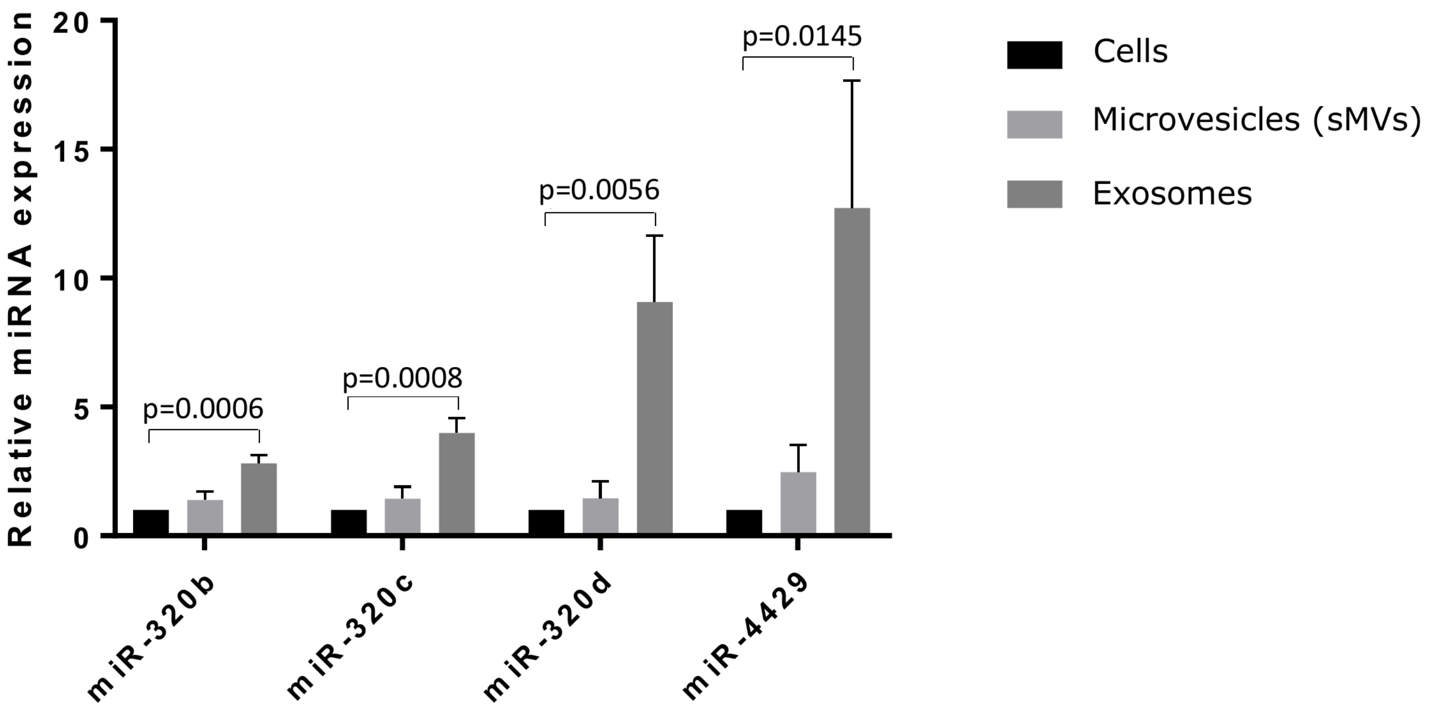

Supplementary Figure 2A

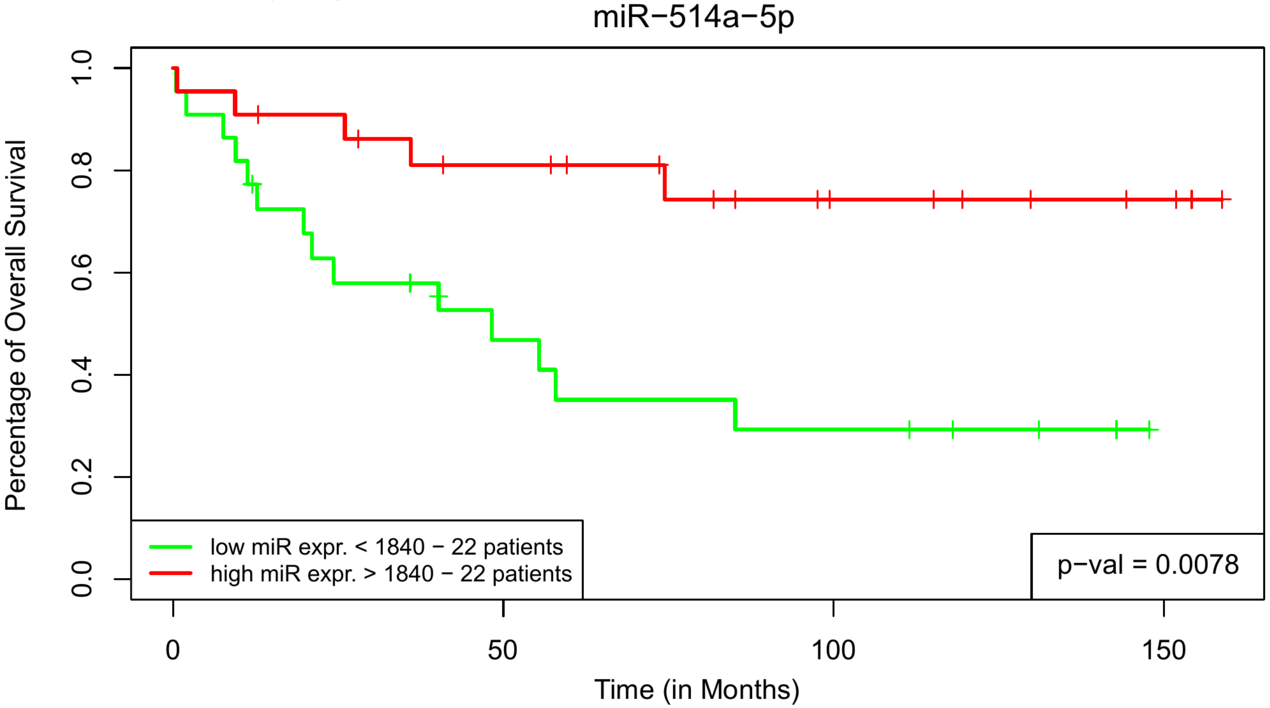


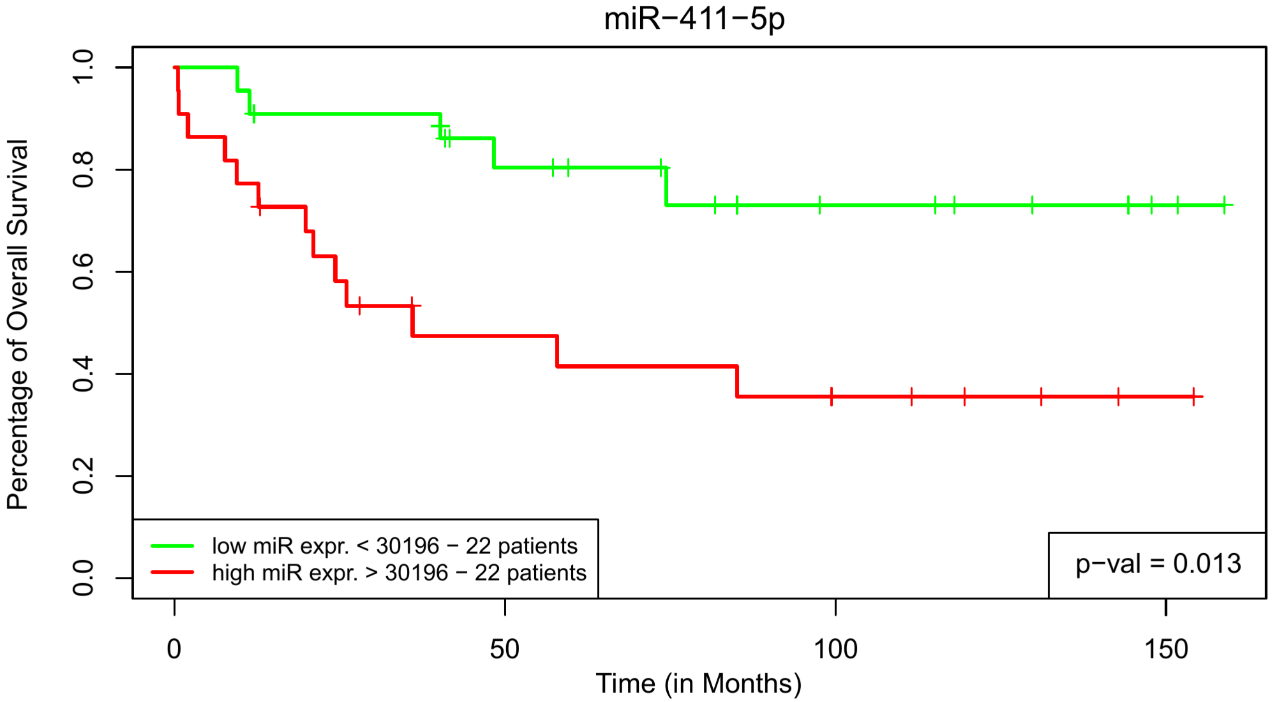

Supplementary Figure 2B

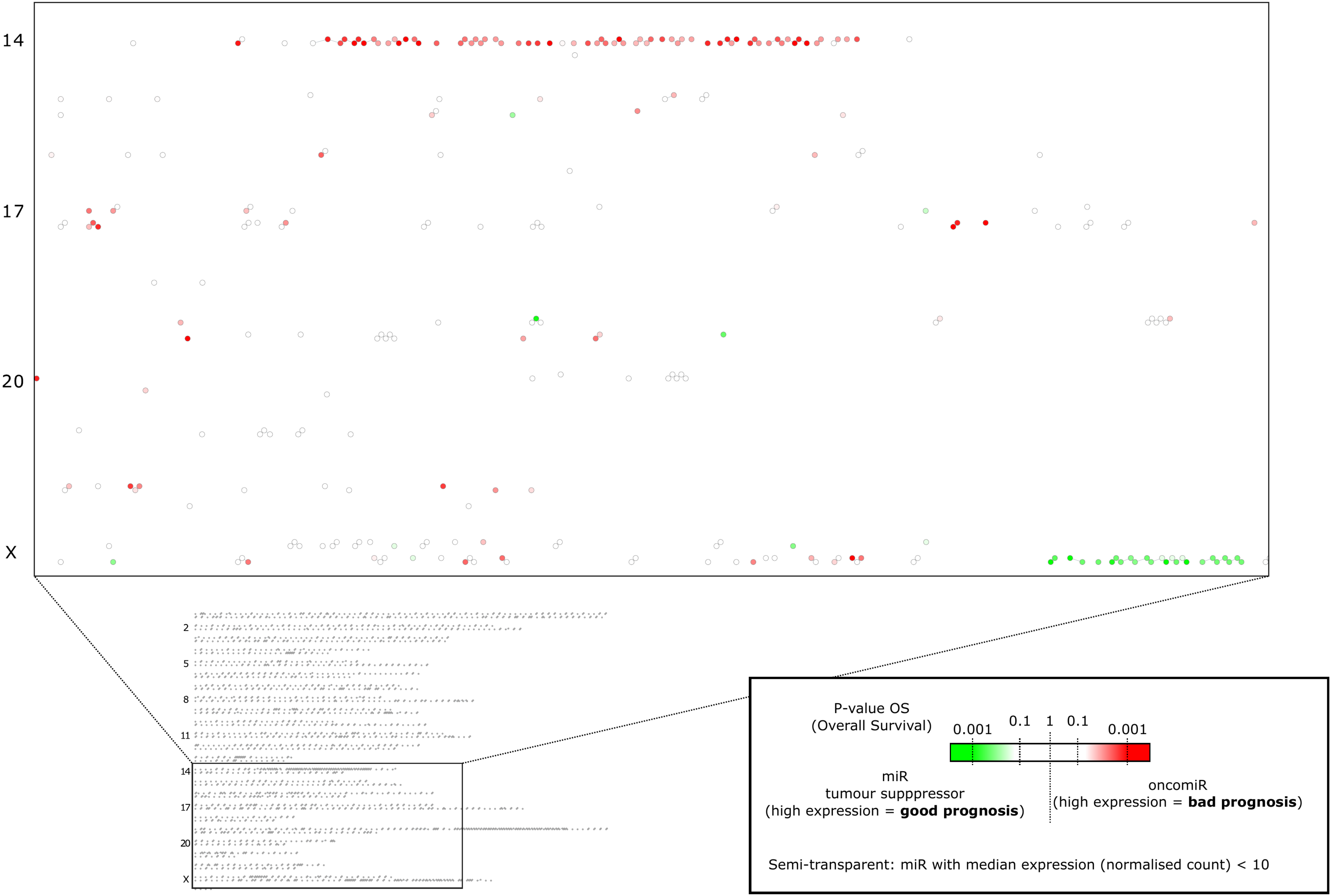

Supplementary Figure 2C

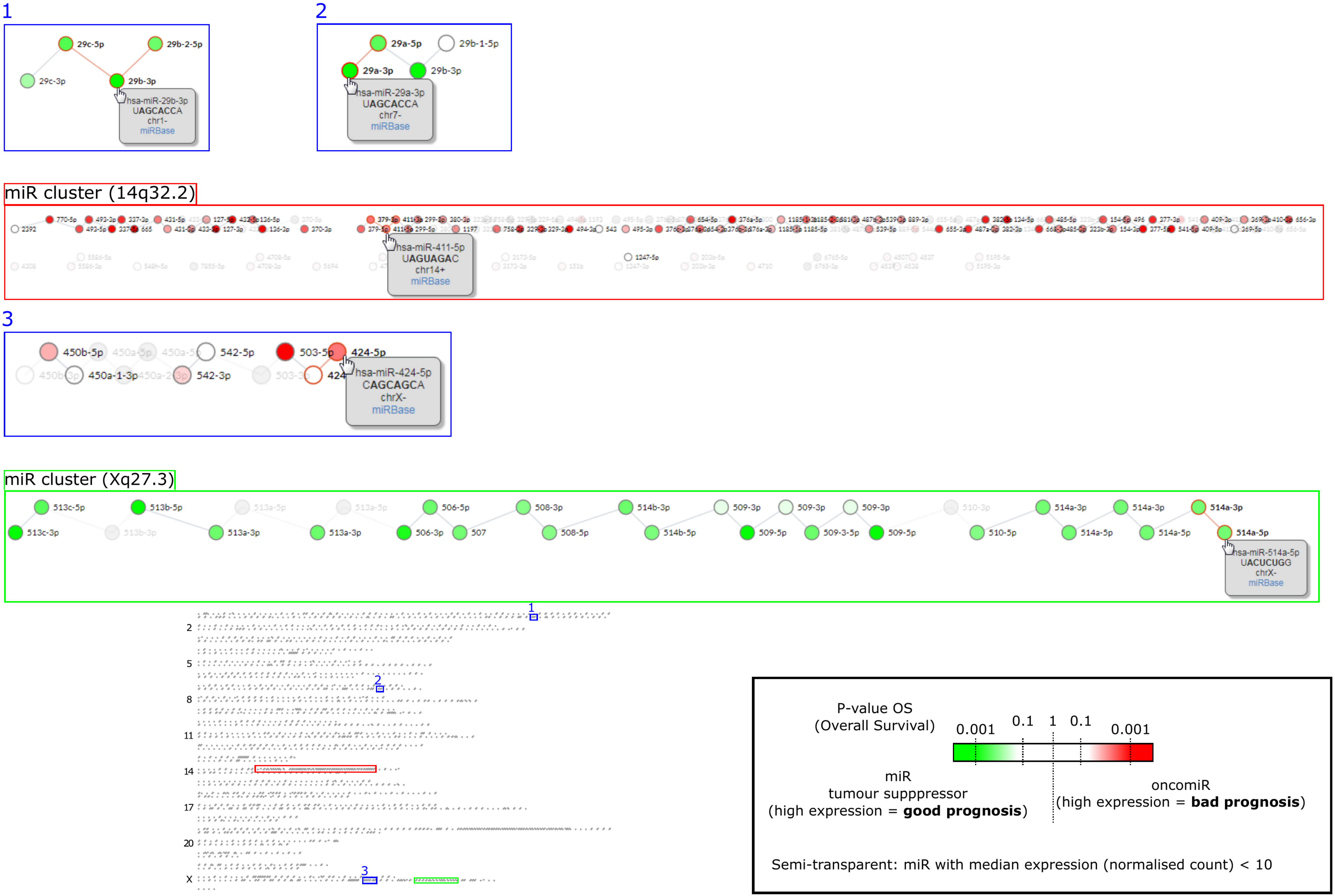

Supplementary Figures 3

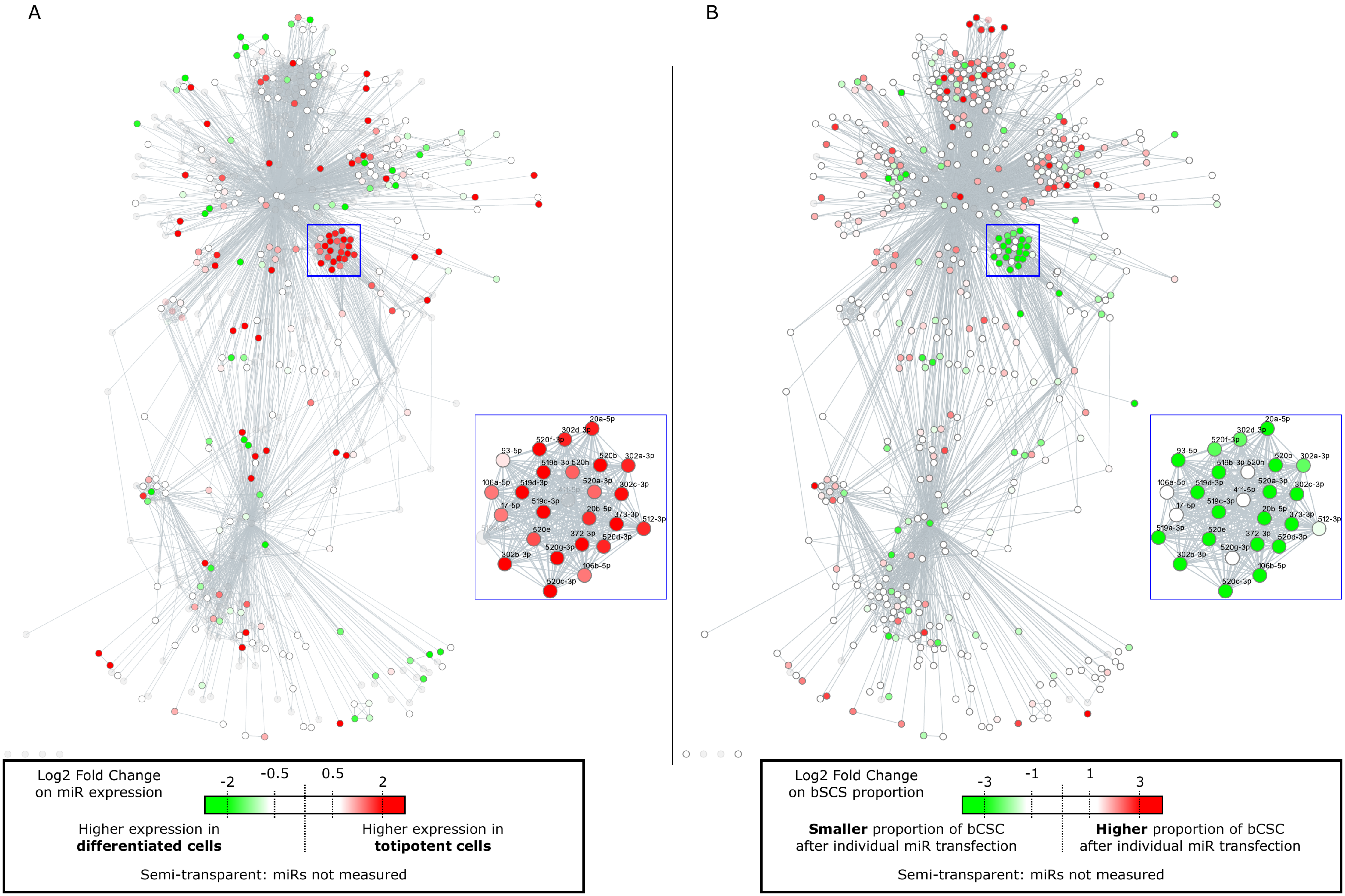

Supplementary Figure 4A

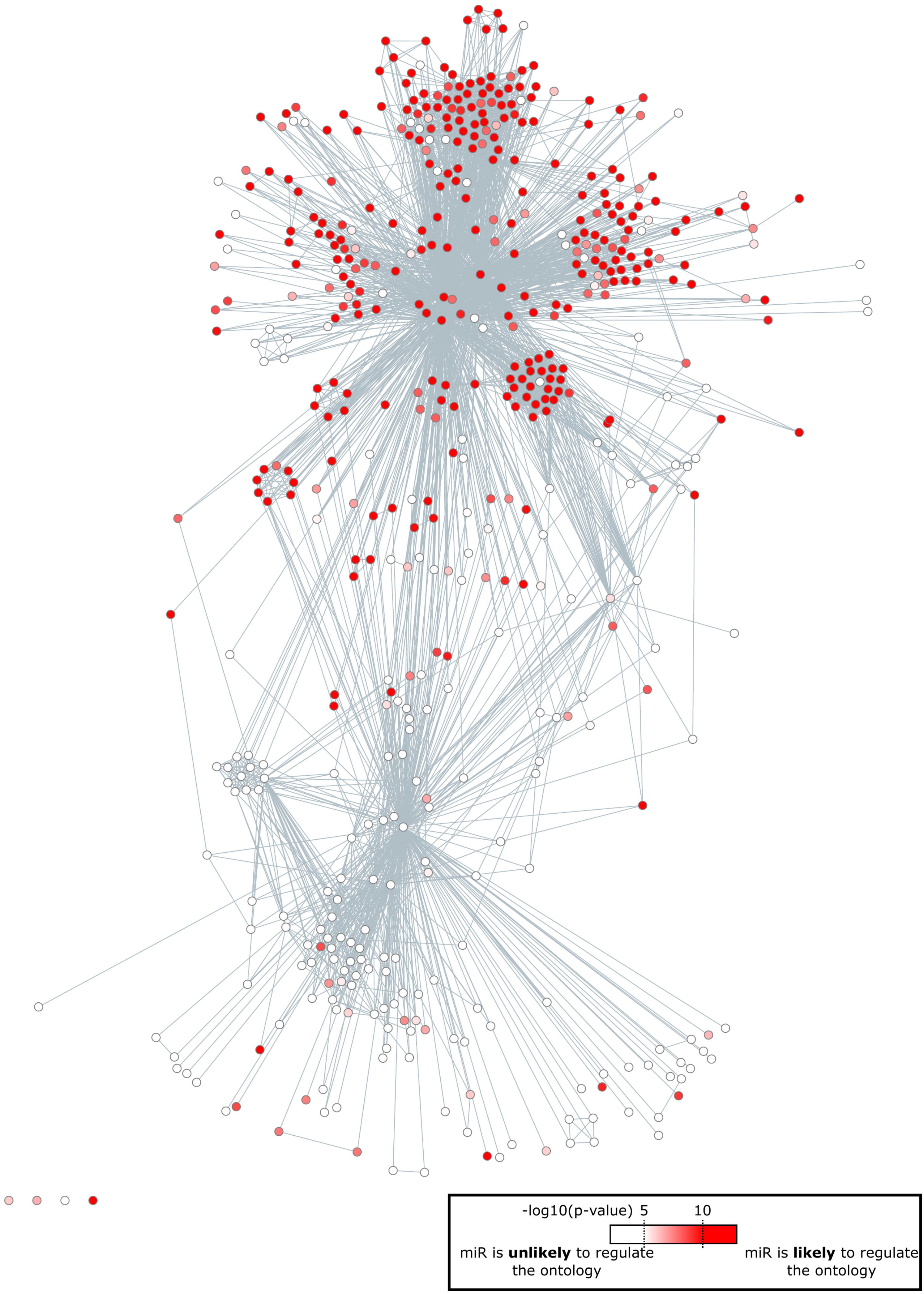

Supplementary Figure 4B

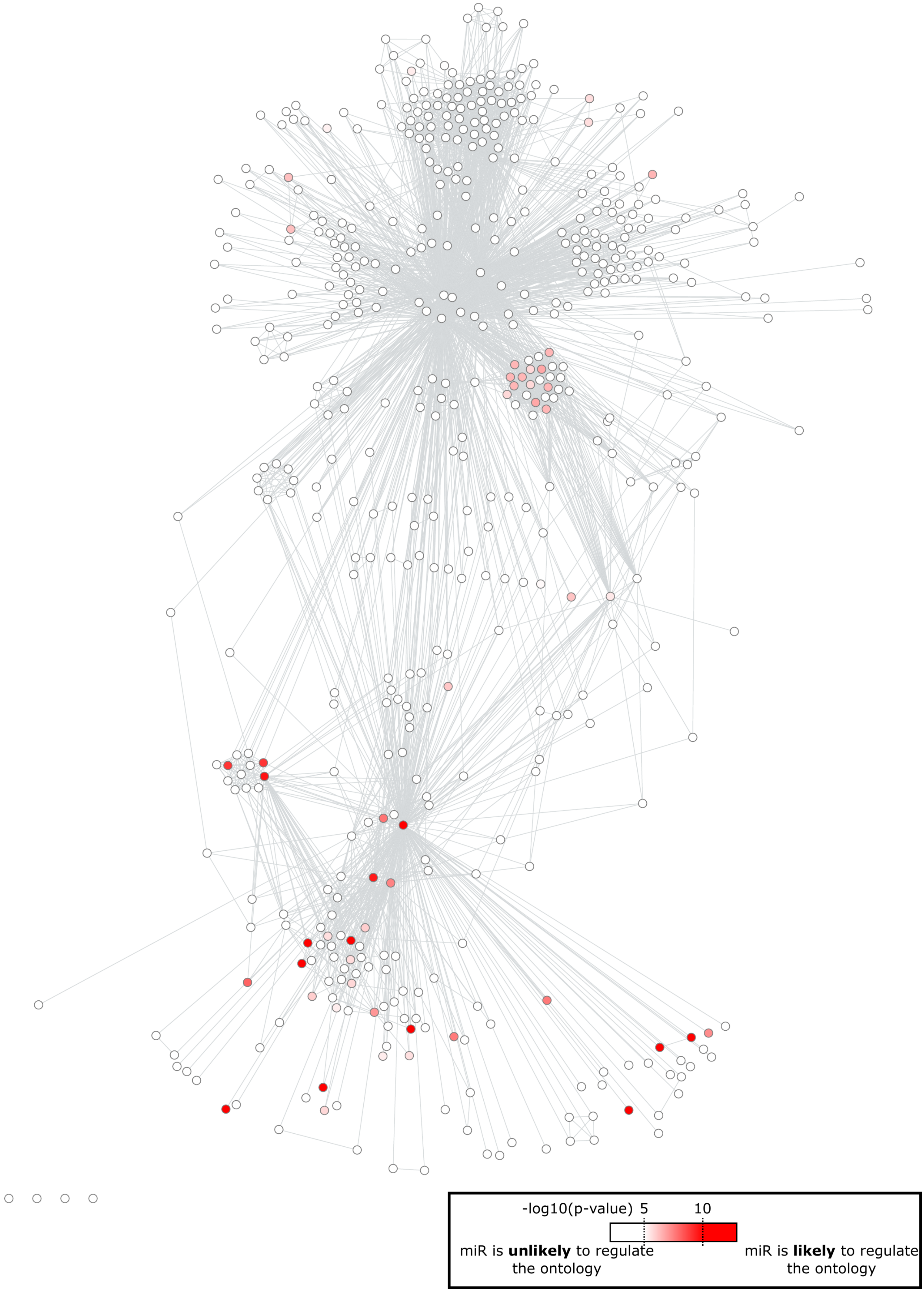

Supplementary Figure 5A

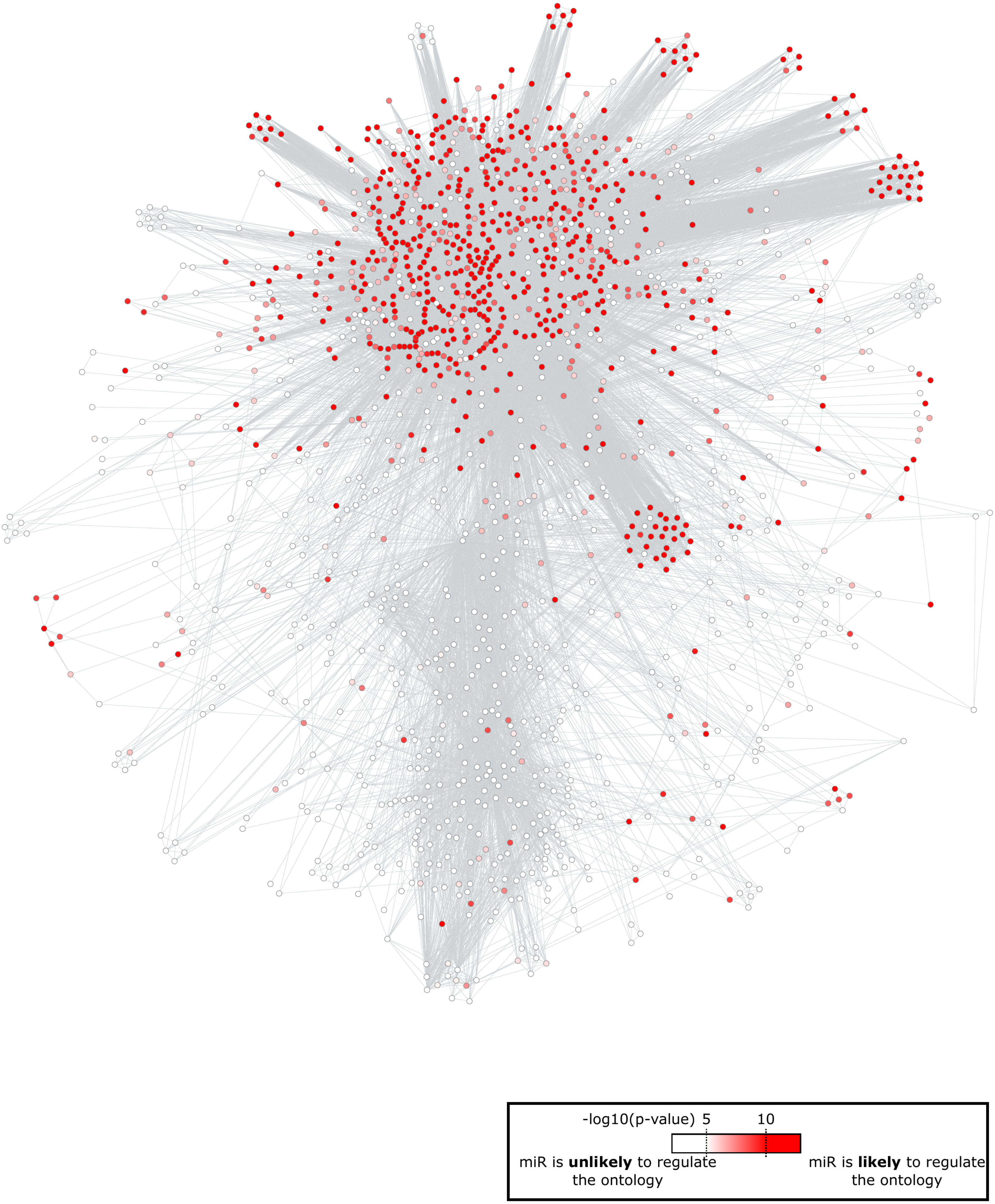

Supplementary Figure 5B

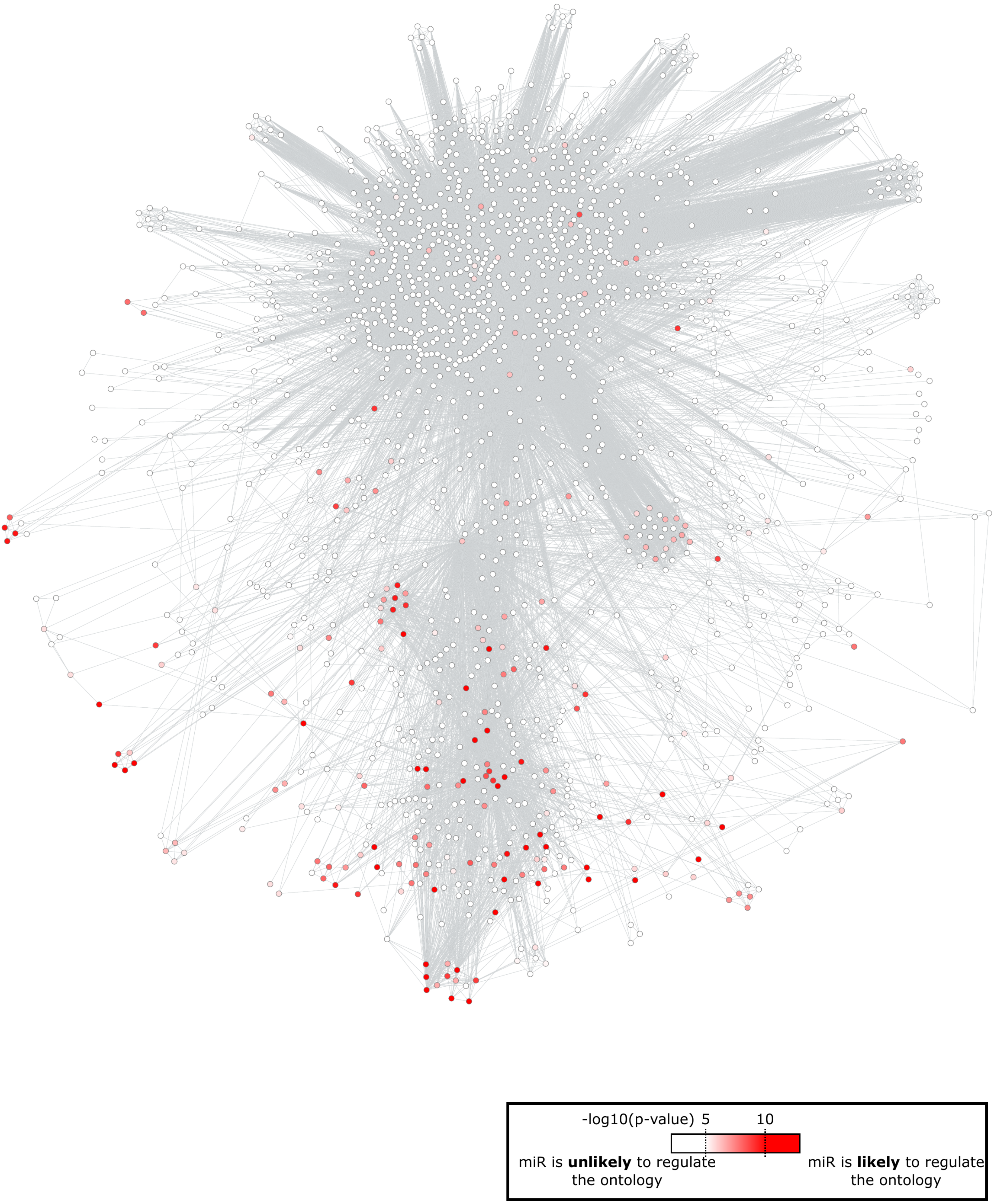
